# Rapid transcriptional response to a dynamic morphogen by time integration

**DOI:** 10.1101/2025.06.09.658715

**Authors:** Susanna E. Brantley, Jacqueline Janssen, Anna Chao, Massimo Vergassola, Shelby A. Blythe, Stefano Di Talia

## Abstract

During development, cells must interpret extracellular signals with speed and accuracy. While morphogen gradients pattern tissues, how cells respond to dynamic morphogens remains unclear. Here, we investigate how dorsal patterning in the *Drosophila* embryo is specified by a rapidly evolving BMP gradient. Using a live reporter of BMP pathway activity and nascent transcription reporters, we find that gene expression is best predicted by time integration of BMP signaling, rather than instantaneous levels. However, in *sog* mutant embryos with broad BMP activity, integration alone fails to predict gene expression outside the normal domain. We show that the transcription factor Zen lowers the signaling threshold required for activation, enabling integration to drive rapid transcriptional responses even at low BMP levels. Together, these results suggest that cells interpret dynamic morphogen signals through the combined action of temporal integration and spatial competence, providing a framework for robust pattern formation on fast developmental timescales.

## Introduction

Many crucial biological decisions, particularly in early development, unfold with great accuracy on rapid timescales, demanding that cells respond to extracellular and intracellular signals with both speed and precision. Morphogen gradients represent a central mechanism controlling developmental patterning [1]. However, most previous models assume that the morphogen input is relatively stable over time. A classic example of how morphogens instruct tissue patterning is anterior-posterior patterning in the *Drosophila* embryo, where Bicoid (Bcd) forms a relatively stable concentration gradient that directs precise stripes of gene expression [2–7]. In this system, positional information is reliably interpreted from a steady-state gradient [6], and many useful models of morphogen-based patterning have built on this ability of cells to read out such steady-state gradients to make cell fate decisions [8]. However, many developmental systems do not operate under steady-state conditions, such as those involving tissue growth or zygotically encoded morphogen gradients [9–11]. Studies of ERK, Shh, and TGF-beta family signaling highlight how signaling history or dynamics changes can be a better predictor of cell fate than a single snapshot of pathway activity [10,12–14]. Still, most of this work has focused on cell fate decisions that evolve over many hours or days. How cells accurately regulate transcription on timescales of minutes remains poorly understood. This question is particularly important during early embryogenesis when patterning decisions unfold within minutes, as well as in adult systems like neurons, where immediate early gene activation occurs soon after synaptic stimulation [17–18]. It remains unclear whether the principles of signal integration established in slowly-evolving systems apply to fast, dynamic signaling environments and how cells interpret signals that are still changing when gene expression decisions need to be made.

Dorsal patterning in the early *Drosophila* embryo offers a powerful system to study rapid input-output responses to dynamic signaling. This patterning is driven by an evolving gradient of BMP activity, which cells must interpret within minutes prior to gastrulation to specify dorsal ectoderm and the extraembryonic tissue of the fly, called the amnioserosa [19–21]. BMP signaling begins during the 14^th^ interphase of embryonic development, 2-3 hours after egg lay, as the syncytial *Drosophila* embryo begins to cellularize prior to gastrulation. During this period, the BMP ligand, *dpp,* is expressed broadly across the dorsal half of the embryo, leading to initially diffuse BMP activity across the dorsal domain. Despite this broad ligand expression, BMP signaling becomes progressively enriched at the dorsal midline by the onset of gastrulation. This refinement is mediated by the secreted BMP inhibitor, *sog*, which is expressed in ventrolateral regions and binds BMP ligands to prevent ligand binding to receptors while also promoting long-range diffusion of ligand complexes [22–24]. These Sog-BMP complexes are released by the dorsally expressed protease, *tld* [20,25]. This source-sink mechanism coupled with positive feedback [26], concentrates BMP signaling activity at the midline. This dynamic evolution of the BMP morphogen gradient was observed over 20 years ago in fixed embryos using antibodies against phosphorylated Mad (pMad), the activated form of the BMP pathway transcription factor [27–28], or the co-SMAD, Medea, which is required for nuclear pMad localization [29]. Since then, genetic experiments and mathematical models have provided strong support for the source-sink inhibitor model of BMP gradient formation [30–32].

Despite these observations of this dynamic signaling patterns and studies of expression of BMP targets [33], there is little understanding of how nuclei in the embryo interpret BMP signaling activity across time and space to produce the nested patterns of BMP responsive genes on the dorsal side of the embryo. This is largely due to the lack of live imaging tools to measure the input-output relationship between BMP activity and target gene expression. Here we developed a live reporter of endogenous BMP activity coupled with MS2-MCP reporters of endogenous target gene transcription to capture this input-output relationship in each nucleus of a living embryo. With these tools, we found that the decision to express BMP target genes is made in response to very small changes in BMP signaling levels and that this response sweeps out from the dorsal midline following a moving threshold of BMP activity. Notably, we find that the integrated BMP signaling response is more predictive of the onset of gene expression rather than an instantaneous threshold. We offer evidence that BMP target genes respond very quickly to low levels of BMP signaling via the catalyzing activity of the transcription factor, Zen. However, morphogen signaling input alone is not sufficient to fully predict gene expression domains. In *sog* mutant embryos with altered BMP dynamics, nuclei outside of the normal spatial expression domain were less likely to express certain BMP target genes than our integrated threshold model would predict. These observations suggest the existence of additional spatially patterned regulatory mechanisms that restrict gene expression even in the presence of sufficient morphogen signaling. Thus, our findings shed light not only on how rapidly changing signals are decoded over time, but also to what extent morphogen inputs alone can account for spatial gene expression patterns during early development.

## Results

### BMP target genes are first expressed in a dynamic signaling environment

To determine when BMP target genes are first expressed relative to BMP signaling activity, we examined carefully staged fixed embryos stained for pSMAD and smFISH against BMP target mRNAs. Embryos were staged by the progression of the cellularization front during interphase of cycle 14 (Fig 1 A-C), allowing us to align developmental time across samples with approximately 10-minute time resolution [34]. From these fixed tissue time course data, we constructed a map of BMP activity dynamics in time and space (Fig 1D, Suppl Fig 1B). These data show that the BMP activity gradient begins to form about 20 minutes after anaphase of nuclear cycle 13, with BMP activity increasing most rapidly at the midline. As predicted, the BMP activity gradient increases more uniformly across the dorsal side of the embryo in *sog* mutant embryos, resulting in a shallower and wider gradient (Fig 1E, Suppl Fig 1C-D). We also quantified spatially averaged mRNA expression for several BMP target genes during nuclear cycle 14, prior to gastrulation (Fig 1F-H). Specifically, we chose known BMP target genes with variably sized expression domains [35–37]. We computed the input-output relationship between nuclear pSMAD levels and the probability of transcription in nuclei grouped in time and space. We found that genes expressed more broadly across the dorsal side of the embryo, such as *ush* and *doc2*, were activated at very low levels of BMP activity, and could be detected as early as 30 minutes into nuclear cycle 14 (Fig 1F-G, I-J).

**Figure 1:**
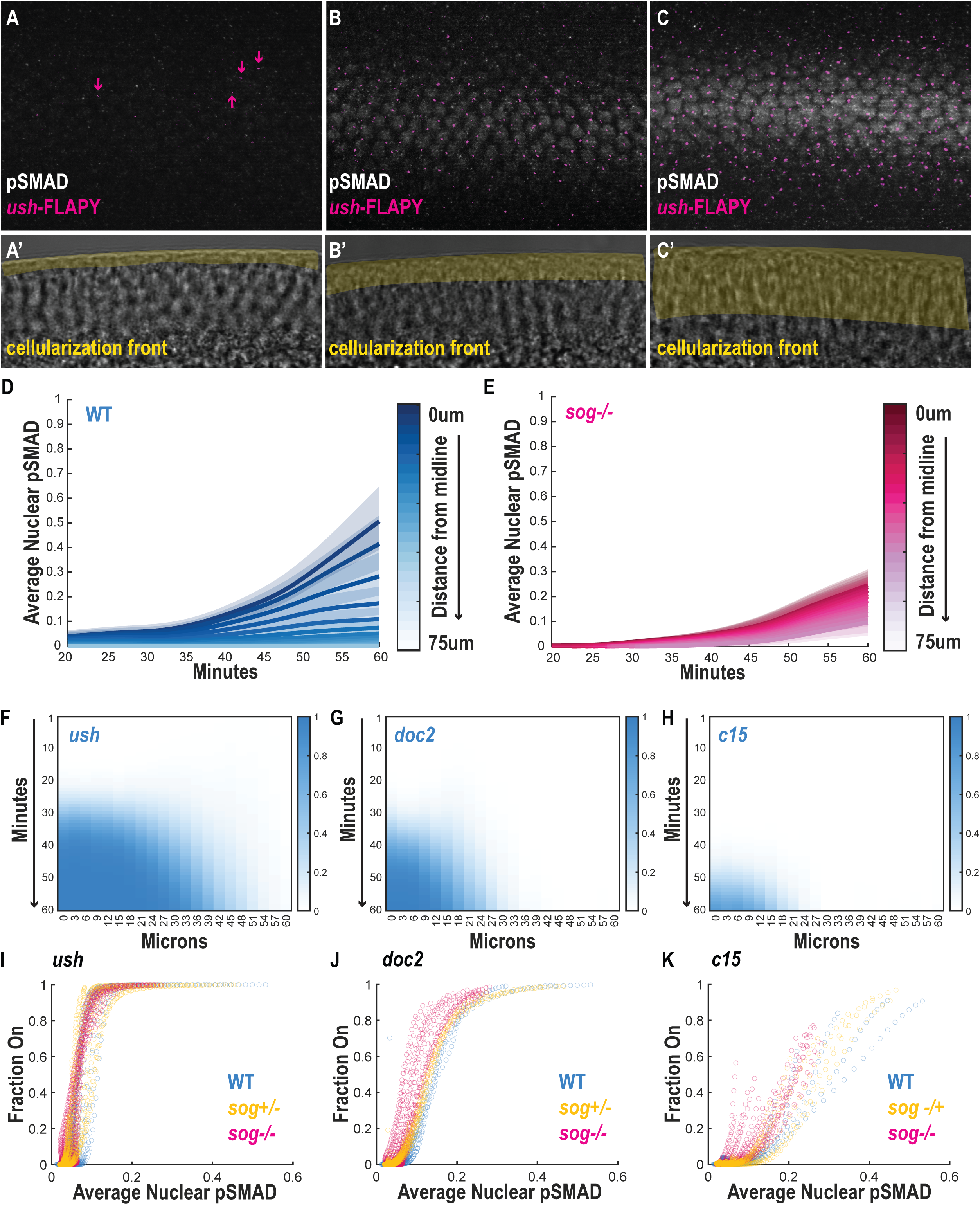
BMP target genes are first expressed concurrently with BMP gradient formation. A-C) Representative fixed wild-type embryos at progressive stages of nuclear cycle 14, staged by the extent of the cellularization front (yellow, A’-C’). Embryos were co-stained for phosphorylated SMAD (anti-pSMAD, grey) and the BMP target gene *ush* mRNA (smFISH, magenta). D) Time-resolved, spatially binned quantification of average nuclear pSMAD across the dorsal-ventral axis in wild type (D) and *sog^S6^* (E) embryos, aligned to developmental time by cellularization front progression F-H) Spatial and temporal quantification of nuclear transcriptional activity (fraction of nuclei with smFISH signal binned by microns from the midline) for three BMP target genes: *ush* (F, n=52 embryos), *doc2* (G, n=32 embryos), and *c15* (H, n=75 embryos). **I–K)** Input-output relationship between average nuclear pSMAD and the probability (fraction on) of target gene transcription for *ush* (I), *doc2* (J), and *c15* (K), in wild-type (blue), *sog^S6^* heterozygous (yellow), and *sog ^S6^* mutant (magenta) embryos.

From these fixed tissue data, it is clear that expression of BMP targets occurs within minutes of exposure to low levels of nuclear pSMAD and at a time when the signaling is rapidly changing over time. Thus, measuring the input-output relationship between BMP activity and target gene expression requires a quantitative method that captures signaling and gene expression dynamics with high temporal resolution. This is difficult to achieve using fixed embryos during the first 30minutes of nuclear cycle 14, as the cellularization front moves slowly at that time (Suppl Fig 1A), making staging imprecise. In addition, we found that the total amount of nuclear pSMAD at a given time did not correlate well with the probability of gene expression for genes like *c15*, which is expressed later and in a narrower spatial domain than the other target genes we analyzed (Fig 1K). Thus, we hypothesized that expression of BMP target genes might not depend on the instantaneous level of signaling but instead might be controlled by other features, such as signaling history. To test this, we developed a live imaging approach to visualize both BMP activity and activation of BMP target gene expression.

### A live reporter of endogenous BMP signaling

In the *Drosophila* embryo, BMP signaling ultimately promotes the nuclear translocation of a SMAD complex, Mad/Medea, which activates transcription (Fig. 2A) [29,38]. We took advantage of this mechanism of action to develop an endogenous reporter of BMP activity. Tagging the C-terminus of the co-SMAD Medea with eGFP or TagRFP resulted in homozygous-viable flies that we used to measure pathway activity in the fly embryo. Immunostaining using a pSMAD antibody has been used historically in the field to visualize and quantify BMP activity. We found a strong correlation between the nuclear levels of GFP-Med and pSMAD antibody staining (Suppl Fig 2A). We also performed ChIP-seq using an anti-GFP antibody on cycle 14 embryos expressing GFP-Med and an anti-pSMAD antibody on *w^1118^* embryos. We compared the top peaks from these datasets to each other and removed any peaks found in a control anti-GFP ChIP in *w^1118^* embryos. A majority of high quality GFP-Med and pSMAD ChIP peaks overlap, providing further evidence that GFP-Med is a functional protein (Suppl Fig 2B-D).

**Figure 2:**
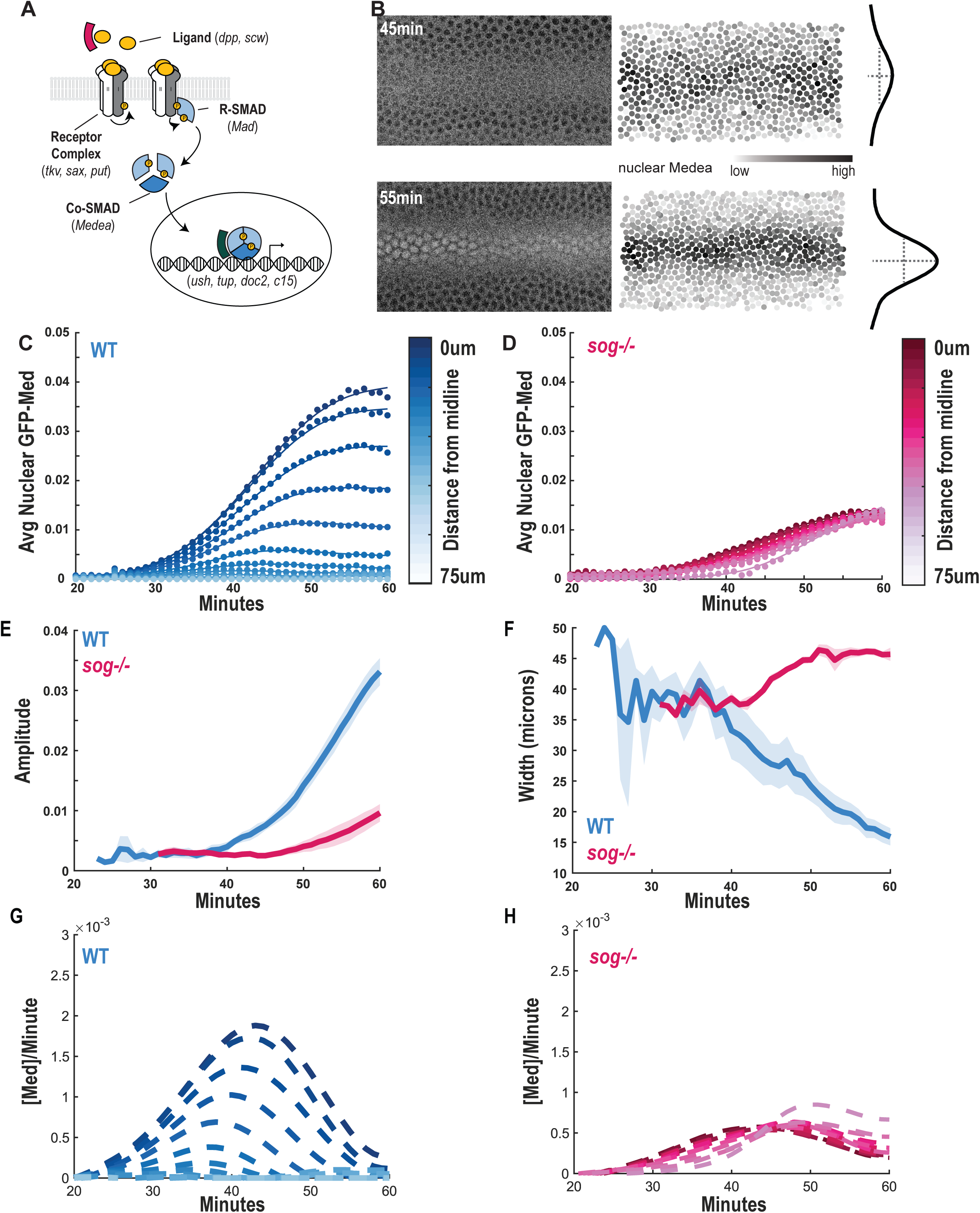
A live imaging reporter reveals dynamic BMP signaling activity. **A)** Schematic of the Drosophila BMP signaling pathway. BMP ligands (Dpp and Scw) bind to a receptor complex (Tkv, Sax, and Put), leading to phosphorylation and nuclear translocation of SMAD proteins (Mad and Medea), which activate transcription of target genes. GFP-Med reporter lines were generated by endogenously tagging the co-SMAD Medea on the protein’s C-terminus by CRISPR/Cas9 mediated integration. **B)** Example of gradient quantification from a live embryo expressing GFP-Medea. Nuclei were segmented via His-iRFP (not shown), and BMP signaling was measured as from nuclear GFP-Medea levels (color-coded, darker shades indicate higher signal). BMP gradients were fit along the dorsal-ventral axis at each time point in nuclear cycle 14 to extract amplitude and width at half-maximum. **C–D)** Time-resolved dynamics of nuclear GFP-Medea accumulation in wild-type (C, n=8) and *sog* mutant (D, n=3) embryos. Data shown are average nuclear intensity in 5 µm spatial bins from the dorsal midline, with 1-minute temporal resolution. **E–F)** Temporal evolution of BMP gradient amplitude (E) and width at half-maximum (F) in wild-type and *sog* mutant embryos. **G–H)** Rate of change in GFP-Medea levels across the dorsal-ventral axis in wild-type (G) and *sog* mutant (H) embryos. In wild-type embryos, the BMP gradient grows most rapidly at the midline; in *sog* mutants, the gradient evolves more uniformly across space.

To observe BMP signaling on a per-nuclear basis over time (Fig 2B), we imaged fly embryos carrying the GFP-Med reporter and a fluorescent nuclear marker (HisiRFP) [39] from cell cycle 13 to the onset of gastrulation at minute-by-minute resolution. Nuclear GFP-Med concentration began to increase across the dorsal side of the embryo about 30min after anaphase of cell cycle 13 and continued at the dorsal midline until gastrulation, observed as a decrease in width and an increase in amplitude of the signal over time (Fig 2C, 2E-F). As predicted from our fixed imaging data, BMP signaling shows differences in both absolute level and rate of change across the dorsal-ventral axis of the embryo (Fig 2C, 2G). BMP signaling at the midline increases at a faster rate and does not reach a plateau before morphogenesis begins 60min into cell cycle 14, while BMP activity in nuclei 10 μm or more from the midline (about 2 cell diameters) increases more slowly and plateaus prior to gastrulation. In contrast, using this BMP sensor in a *sog* mutant background, we observed less difference in dynamics at the midline compared to lateral regions of the embryo (Fig 2D, 2H). This results in lateral regions of the embryo experiencing higher levels of BMP signaling than in a wild-type embryo while the midline experiences lower BMP signaling. BMP signaling dynamics in *sog* heterozyogous embryos showed an intermediate phenotype (Suppl Fig 2G-H). These observations have been previously reported in fixed tissues [40], further confirming the utility of GFP-Med as a readout of BMP activity and the use of *sog* mutants as a tool to modulate pathway activity.

### Input-output dynamics at BMP target genes

To monitor how the activation of validated downstream targets is influenced by BMP signaling levels, we combined our BMP reporter with MS2 reporters that allow visualization of nascent transcripts. Using an MS2 reporter of *ush* mRNA expression [33], we could correlate the amount of BMP activity with onset of target gene transcription (Fig 3A). The *ush*-MS2 reporter was first detected in cells at the midline at around 35 minutes after anaphase of nuclear cycle 13, while transcriptional onset time is delayed in nuclei further away (Fig 3B), consistent with previous reports using this *ush*-MS2 reporter and our fixed tissue data. Thus, activation of this early BMP target gene is not determined by the position of a cell within a steady-state morphogen gradient. Rather it is responding during a period of fast-evolving signaling dynamics.

**Figure 3:**
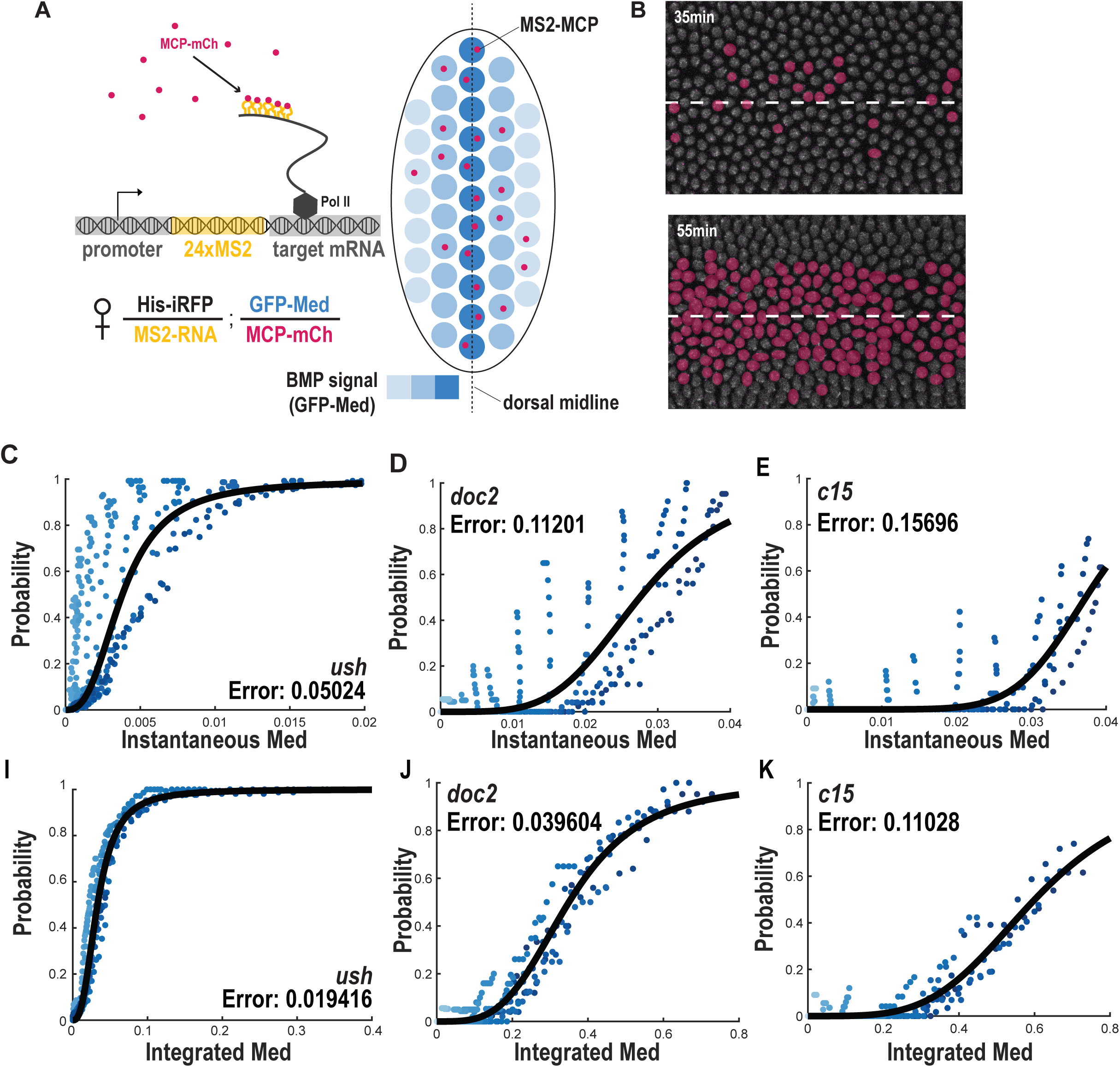
Integrated BMP signaling predicts target gene activation. **A)** Schematic of the live imaging approach to quantify the relationship between BMP signaling dynamics (via nuclear GFP-Medea) and nascent transcription of BMP target genes (via MS2-MCP dot detection). **B)** Example time points showing nuclei with active transcription of *ush* (magenta, MCP-MS2 dots) overlaid on nuclear GFP-Medea signal. Transcriptional onset is first detected at the dorsal midline and progressively later in more lateral nuclei. **C–E)** Relationship between instantaneous nuclear GFP-Medea levels and the probability that a nucleus has initiated transcription for three BMP target genes: *ush* (C, n=8), *doc2* (D, n=2), and *c15* (E, n=2). Each point represents a spatiotemporal bin (5 μm × 1 min). Curves show Hill function fits to the data. Color indicates position from the midline (dark blue indicates the midline, light blue indicates 75 μm away). **F–H)** Delayed-threshold models comparing the probability of transcriptional activation to the instantaneous nuclear GFP-Medea level with a lag time of 5 min (F), 10 min (G), or 15 min (H). Curves show Hill function fits to the data. Color indicates position from the midline (dark blue indicates the midline, light blue indicates 75 μm away). **I–K)** Relationship between transcriptional probability and integrated GFP-Medea signal for *ush* (I), *doc2* (J), and *c15* (K).I-K) Input-output relationship between the probability that BMP target genes *ush* (I), *doc2* (J), or *c15* (K) is expressed and integrated GFP-Med in nuclei binned in time and space from wild type embryos. Data are fit with a Hill curve every 15 microns from the midline, and color indicates position from the midline (dark blue indicates the midline, light blue indicates 75 μm away). These analyses demonstrate that integrated signaling over time provides a more robust predictor of BMP target gene activation than instantaneous signal levels, indicated by the root mean absolute error (RMSE corrected by the average) for each fit.

To test whether different BMP target genes exhibit distinct temporal activation patterns, we developed new MS2 reporters for *doc2* and *c15*, which are expressed in more dorsally restricted domains than *ush* and are essential for proper amnioserosa formation [35–37]. Using these tools, we found that the timing of initial activation varied among target genes, and that genes activated earlier (such as *ush*) ultimately occupy broader spatial domains by the onset of gastrulation than genes activated later (Fig 1F-H). This suggests that the time of gene activation influences the final width of expression domains and supports a model in which activation thresholds and timing are linked.

We observed that once a BMP target gene promoter became active, it usually remained active and in a bursting regime for the remainder of nc14 before gastrulation (Suppl Fig 3A-C). Thus, we focused on time to initiation of transcription for further analysis. To analyze input-output relationship between signaling and transcription activation, we binned our live imaging data into 5 micron spatial bins (relative to the dorsal midline) and 1 minute time intervals. For each spatiotemporal bin, we computed the following metrics: fraction of nuclei on (i.e. transcribing the gene of interest), instantaneous signal, and integrated signal. The fraction of nuclei on in each spatiotemporal bin was defined as the total number of nuclei which had begun bursting divided by the number of nuclei in the bin. Instantaneous signal was defined as the average nuclear GFP-Med intensity within a spatial bin at each timepoint. Finally, integrated signaling was defined as the cumulative integral of GFP-Med over time for each spatial bin.

With these measurements, we could then infer the relationship between transcription probability and either instantaneous or integrated signaling levels. To formalize this relationship, we modeled the data using Hill functions, which are commonly used to describe sigmoidal responses in biological contexts. In this case, the input is pMad/Medea concentration (*[M]*), and the output is represented by the probability (*P_RNA_*) that a BMP responsive promoter is actively transcribing mRNA. Gene activation is described by a dissociation constant (*K_a_*) and the Hill coefficient (*n*) via the following relationship:

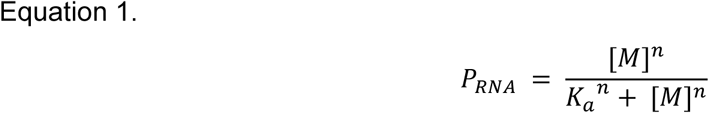

If we plot the probability that a nucleus has begun bursting versus the amount of GFP-Med in the nucleus for spatial-temporal nuclear bins, we find that there is a weak correlation described by a non-linear function (Fig 3 C-E). If we instead compare signaling levels integrated over time with probability that a nucleus has begun bursting, we see a much stronger correlation (Fig 3I-K), suggesting that integrated signaling is a better predictor of gene expression than instantaneous amounts of transcription factor in the nucleus.

To strengthen the evidence that gene activation is controlled by time integrated BMP signaling, we tested whether integration predicts gene expression under altered signaling conditions. To this end, we performed experiments in *sog* mutants, which exhibit expanded and flattened BMP gradients and spatially uniform signaling dynamics (Fig 1E, Fig 2D) [22]. Adding a mutant allele into the live imaging system was genetically challenging, so we turned to a fixed tissue time course of RNA expression, which begins after we can easily stage embryos by the cellularization front. We then mapped binned smFISH data onto live movies of GFP-Med activity from *sog* mutant embryos, allowing us to reconstruct integrated signaling histories for each spatiotemporal bin. If BMP target genes were activated based solely on time integrated signaling levels, we would expect their expression domains to expand in parallel with the broadened gradient in *sog* mutants. Consistent with this prediction, we found that the spatial and temporal expression domains of target genes were indeed expanded in *sog* mutants and heterozygotes (Fig 4A-B, Suppl Fig 3D-I). However, a more detailed analysis revealed a striking breakdown of the integrated input-output relationship in more lateral regions of the embryo (> 45 microns from the midline). Outside of the wild type expression domain, nuclei reached integrated BMP levels comparable to those sufficient to activate transcription in wild type nuclei near the midline, yet they failed to express the target gene at the predicted probability (Fig 4C-D). We fit Hill curves to the integrated signaling data across spatial bins and extracted *K_a_* values from those fits to assess whether different regions of the embryo required different amounts of integrated signal to activate transcription (Fig. 4C-D). This reveals a spatial shift in the activation threshold *K_a_* in regions further from the midline, indicating that the amount of BMP signal needed for activation was increased relative to regions within the wild type expression domain (Fig 4E-H). These findings suggest that while time-integrated BMP signaling is a key determinant of transcriptional activation, it is not sufficient to fully explain the spatial patterning of gene expression. Additional regulatory mechanisms likely constrain target gene activation outside of their normal expression domains.

**Figure 4.**
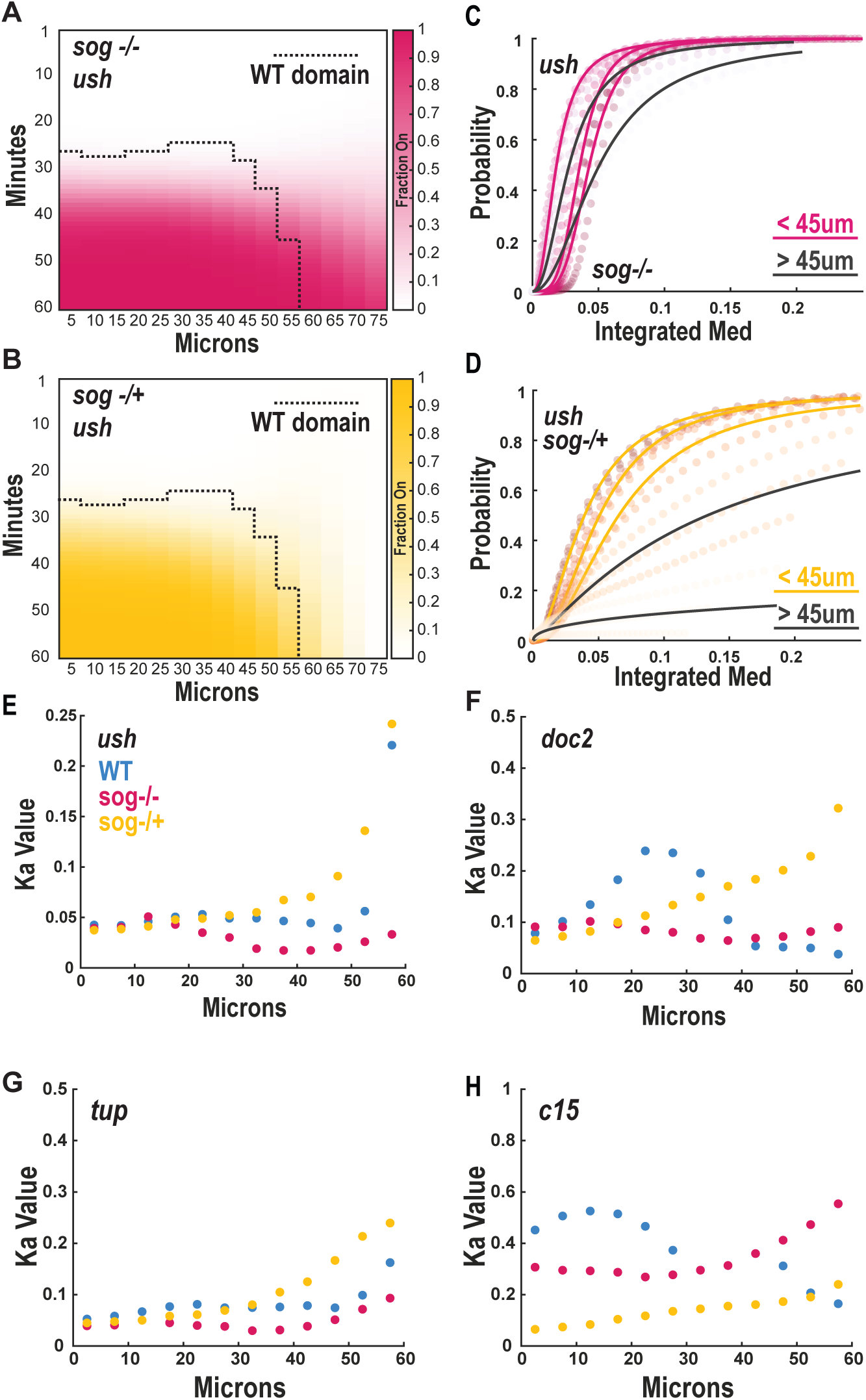
Integration of BMP signaling over time partially predicts target gene expression in *sog* mutants. **A–B)** Spatial and temporal expression patterns of BMP target genes *ush* in fixed embryos from *sog* mutants (A) and *sog* heterozygotes (B), visualized by smFISH. Embryos were staged by the progression of the cellularization front. The wild type expression domain (at least 10% of nuclei on) is outlined with a dotted line. **C–D)** Input-output relationship between integrated GFP-Medea levels (from live imaging data) and the probability of transcriptional activation for *ush* in (C, n=29) *sog* mutants and (D, n=17) heterozygotes. Each point represents a 5 μm × 1 min bin. Data are color-coded by distance from the dorsal midline. Hill curves were fit to collections of 15 μm spatial bins relative to the dorsal midline. While bins close to the midline (yellow or magenta) recapitulate wild-type behavior, lateral bins (>45 μm, grey) deviate from the expected input-output relationship. **E–H)** Spatial distribution of Hill function dissociation constants (Ka) extracted from fits such as those in (C–D), plotted as a function of distance from the dorsal midline. Comparison of Ka values across genotypes (WT-blue, *sog* het-yellow, *sog* mutant-magenta) for *ush* (E), *doc2* (F), *tup* (G), and *c15* (H) highlight the spatial shift in activation threshold with increasing distance from the midline, especially in mutant backgrounds. These data show that while integration of BMP signaling over time explains much of the transcriptional response, lateral nuclei in *sog* mutants fail to activate target genes despite sufficient cumulative signal, revealing additional spatial constraints on BMP target gene expression.

### Modulation of the input-output response to BMP

To gain further insights on how cells respond to integrated BMP signaling, we focused on the role of Zen which is thought to be involved in feed-forward signaling at some BMP target genes [36,41]. To this end, we analyzed *zen^7^* mutant embryos (Fig 5A-C), which have increased levels of BMP signaling [42]. Like the method described for *sog* mutants, we measured the fraction of nuclei in which we could detect a spot of mRNA transcription by smFISH in spatial-temporal bins and mapped these bins onto live imaging data of BMP gradient dynamics from their respective genotypes.

**Figure 5:**
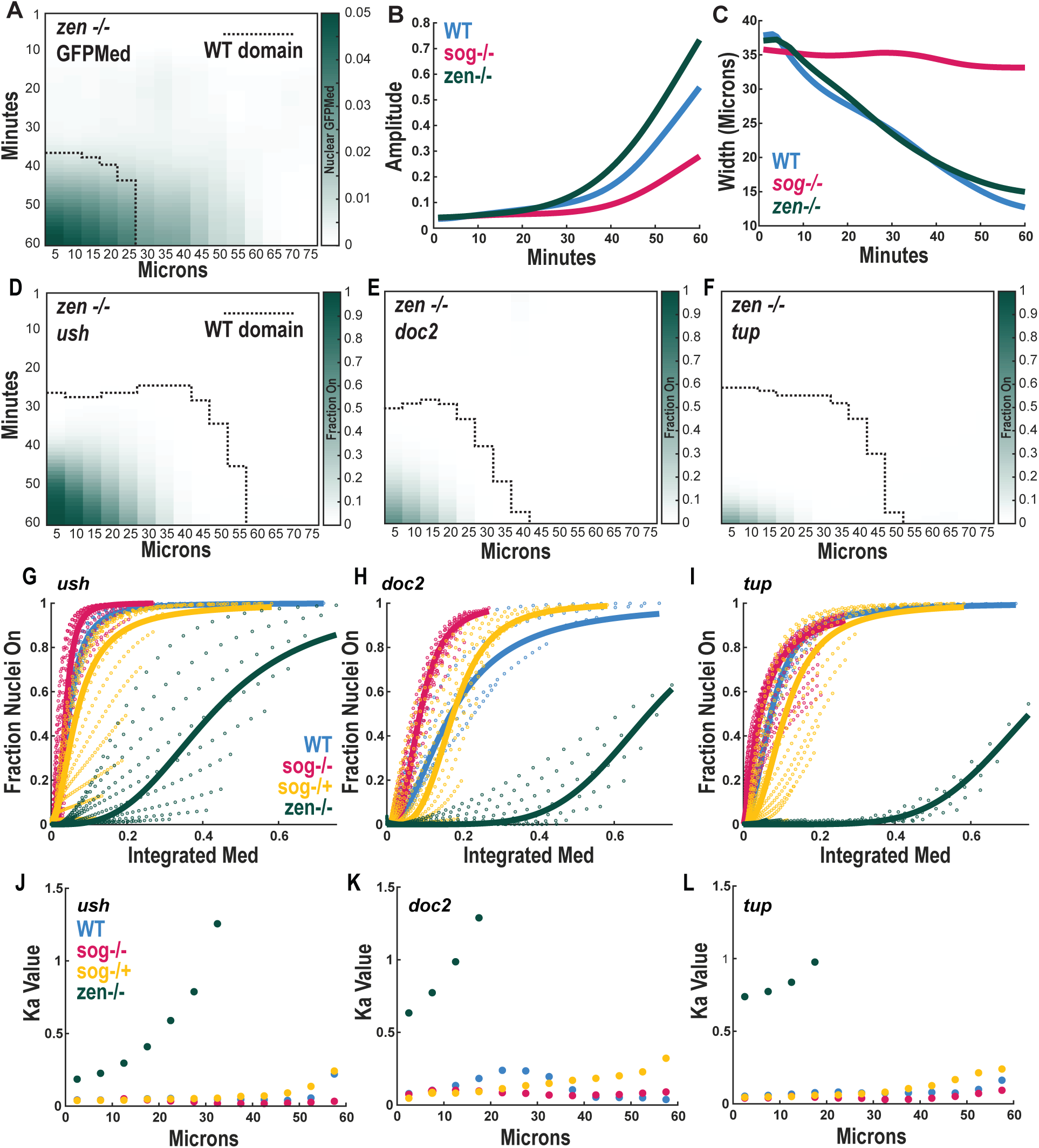
Zen is required for the rapid transcriptional response to BMP signaling. **A)** Spatiotemporal map of nuclear GFP-Medea accumulation in live *zen* mutant embryos (n=2), binned every 5 μm and 1 minute. The dotted line marks the spatial extent of the BMP signaling domain (GFP-Med > 0.01) in wild-type embryos for comparison. **B–C)** Quantification of BMP gradient amplitude (B) and width at half-maximum (C) from fixed wild-type (n=100), *sog* mutant (n=200), and *zen* (n=171) mutant embryos. **D–F)** Spatiotemporal maps of nascent transcription for three BMP target genes (*ush* n=35, *doc2* n=32, *tup* n=33) in *zen* mutant embryos, measured by smFISH. Dotted lines indicate the wild-type expression domain boundary (>10% of nuclei active). **G–I)** Input-output relationships between integrated GFP-Medea levels and the probability of transcription for *ush* (G), *doc2* (H), and *tup* (I), comparing wild-type (blue), *sog* heterozygotes (yellow), *sog* mutants (magenta), and *zen* mutants (green). Data are fit with Hill curves. **J–L)** Dissociation constants (Ka) extracted from Hill fits for each 5 μm spatial bin plotted as a function of distance from the dorsal midline. *zen* mutants show elevated and spatially variable Ka values, indicating reduced transcriptional sensitivity to BMP signaling.

To better understand the relationship between BMP signaling inputs and output of target gene transcription, we again modeled these data using Hill functions. All BMP targets we measured in *zen* mutant embryos showed a reduced sensitivity to BMP signaling and a delay in transcriptional onset despite having higher levels of BMP activity (Fig 5D-F). When we fit a Hill curve to these data, the Hill coefficients were comparable to wild type and *sog* mutants, but the *K_a_* value changed by at least an order of magnitude (Fig 5J-L). Strikingly, in the absence of *zen*, the threshold of integrated signaling was no longer uniform across the dorsal domain. Instead, each BMP target gene appeared to require a spatially dependent threshold of signaling for activation, suggesting a breakdown of the integration-based decoding mechanism. Together with our findings in *sog* mutants, these results point to a role for Zen in enabling promoter responsiveness across the broader dorsal field. We propose that Zen acts to sensitize BMP target promoters, effectively lowering their activation threshold and allowing them to respond to low time-integrated levels of signaling. This has two quantitative effects on the regulation of gene expression. First, it allows the cells on the dorsal side to become sensitive solely to the integrated levels of BMP signaling. Second, it requires cells in lateral regions of the embryo overcome additional positional constraints on gene activation in response to BMP signaling.

It has been previously proposed that Zen acts in a feed-forward loop to activate genes, such as *c15* [36,41]. This hypothesis relies on the fact that *zen* itself becomes a BMP target gene during nc14. This can be seen in the dynamics of *zen* mRNA (Fig 6A). *zen* is first expressed in the cleavage cycles under the control of Zelda, and early in nc14 we observed that *zen* mRNA is actively transcribed in a broad pattern across the entire dorsal side of the embryo. Over time, this broad pattern sharpens and *zen* mRNA becomes more limited to the dorsal side, most likely due to its regulation by BMP signaling. However, it is unclear how to reconcile this with our observation that Zen impacts the expression of BMP targets across their entire expression domain, including the most lateral regions. Zen protein levels during cycle 14 had not been previously characterized and it remained unclear how rapidly the protein spatial pattern changes. To resolve this conundrum, we decided to develop new transgenic lines to visualize Zen protein dynamics. Traditional fluorescent protein tags are limited in this context due to their relatively slow maturation time, which fail to report rapid changes in protein levels, especially for zygotic genes like *zen* that are transcribed and translated within minutes. To overcome this limitation, we used the LlamaTag system [43], which allows for real-time visualization of protein dynamics without the need for rapid fluorophore maturation. We generated an endogenous zen-Llama Tag allele, in which a GFP nanobody is fused to the C-terminus of the Zen protein. In embryos expressing low levels of maternally deposited GFP, the nanobody rapidly binds GFP, enabling us to track Zen protein localization in the nucleus with minimal delay (Fig 6B-E). Quantification of Zen protein levels showed that Zen protein increases uniformly on the dorsal side of the embryo over the course of the last 3 cleavage cycles, well before it is under the control of BMP activity (Fig 6F). By the onset of gastrulation, the pattern of Zen protein remains very broad, displaying only a very small decrease along the DV axis, resulting in the width of the gradient being much broader than the BMP gradient (Fig 6G). These observations suggest that the dorsal sharpening of Zen pattern happens after the onset of transcription of BMP targets. Collectively, our data suggest that Zen acts as a general catalyst to BMP target gene activation by lowering the levels of signaling needed for activation and allowing targets to respond to very low levels of signaling in mid-cycle 14.

**Figure 6:**
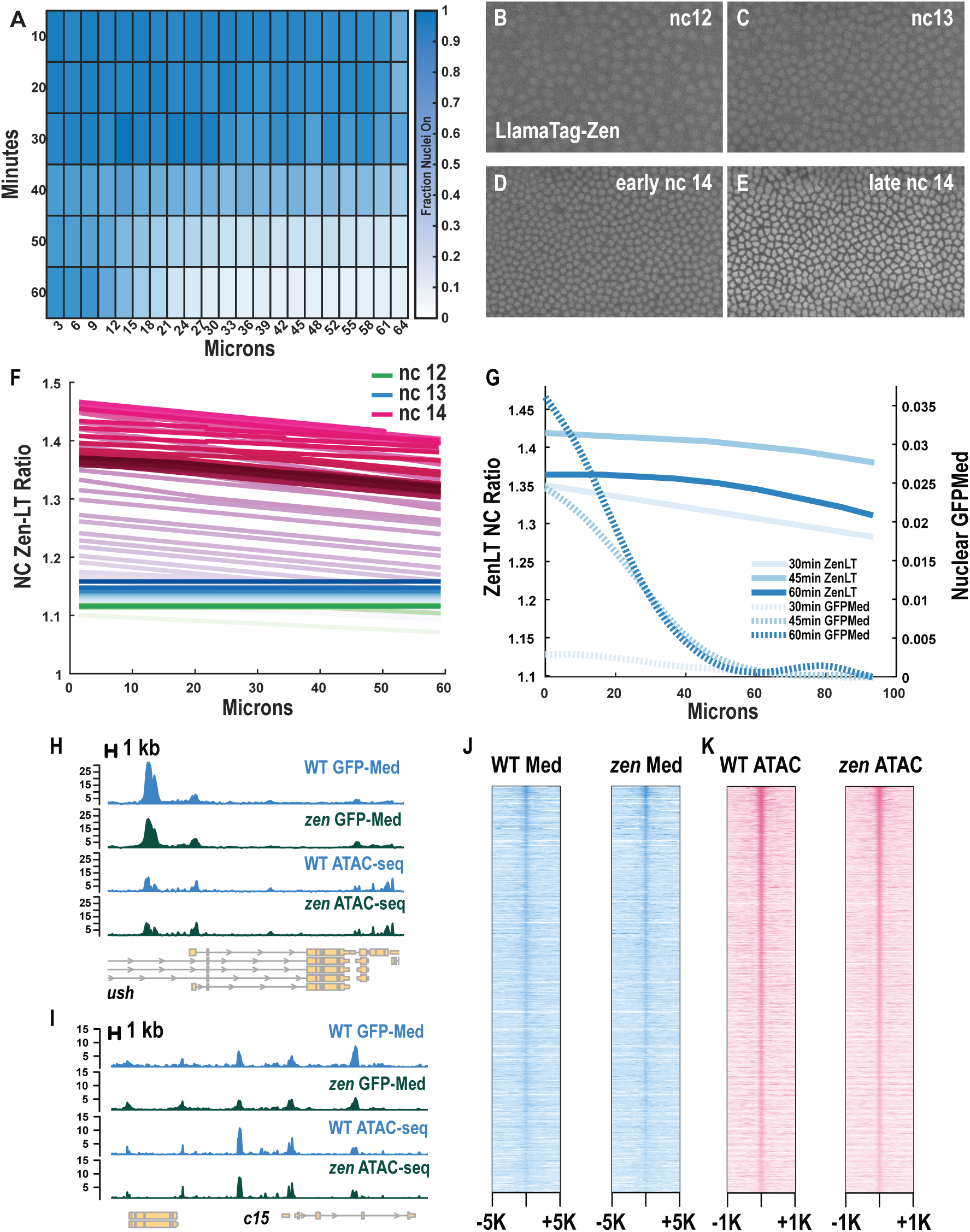
Zen protein is broadly present across the dorsal side during BMP target gene activation. **A)** Spatiotemporal dynamics of *zen* mRNA expression in wild-type embryos, quantified from fixed samples (n=59). Expression is initially broad across the dorsal region and refines over time during nuclear cycle 14. **B–E)** Live imaging of Zen protein using the LlamaTag system in embryos expressing maternally deposited GFP (vasa-GFP), Zen-LT, and His-iRFP (not shown). Representative frames from successive nuclear cycles show nuclear accumulation of Zen-LT prior to gastrulation. **F)** Quantification of the nuclear-to-cytoplasmic ratio of Zen-LT over three nuclear cycles, showing a steady increase in nuclear Zen protein levels prior to and during early nc14 (n=3). **G)** Comparison of the spatial width of the Zen protein gradient and the BMP (GFP-Med) gradient across nc14. Zen protein remains broadly distributed compared to the sharper BMP signaling gradient. **H–I)** ChIP-seq (anti-GFP for GFP-Med) and ATAC-seq tracks at two representative BMP target genes in wild-type (blue) and *zen* mutant (green) cycle 14 embryos. **J)** Summary of GFP-Med ChIP-seq peak locations and intensities in wild-type and *zen* mutant embryos, showing consistent SMAD binding across genotypes. **K)** Summary of ATAC-seq peaks in wild-type and *zen* mutant embryos, indicating no significant change in chromatin accessibility at BMP target loci.

To investigate how Zen might be promoting transcription at BMP target gene promoters, we assayed chromatin accessibility and SMAD binding in *zen* mutant embryos. We performed single-embryo ATAC-seq in wild type and homozygous mutant *zen^7^* embryos staged approximately 15min into cell cycle 14. Nearly all ATAC peaks were consistent between wild type and *zen^7^* mutant embryos, including ATAC peaks near BMP target genes, suggesting that the slower and lower gene expression observed in *zen^7^* mutants is not due to a lack of access to the chromatin (Fig 6H-I, 6K). We also performed ChIP-seq in wild type and *zen^7^* mutant GFP-Med embryos using a GFP antibody and found that *zen* is also not required for Medea binding at BMP target genes (Fig 6H-I, 6J). Thus, we suggest that Zen directly binds these genes to facilitate gene activation and recruitment of transcriptional machinery.

## Discussion

### Live imaging reveals dynamic reshaping of the BMP gradient

Reaction-diffusion models of BMP signaling dynamics have been used to predict the evolution of gradient shape over time. These predictions suggested that the gradient should evolve rapidly following the production of BMP ligands, such that a steep gradient is observed after 30min of simulation [24,32]. However, these models were inconclusive as to whether the gradient became wider or narrower over time and how quickly the amplitude reached its peak. Direct observation of signaling dynamics shows that the gradient narrows over time because the rate and persistence of an increase in signaling is a function of position relative to the dorsal midline. Previous models of BMP dynamics have focused on modeling the binding of ligand to receptor, rather than activity of the SMAD transcription factor, with the assumption that there is a fast, linear relationship between the two. Here, we revisited this model with the goal of using dynamic live imaging to explore the input-output relationship between SMAD activity and target gene transcription.

To this end, we developed a powerful new tool for visualizing BMP signaling dynamics in the early fly embryo. This tool faithfully reports wild type BMP activity, as flies that express GFP-Med are homozygous viable, and ChIP-seq peaks for GFP-Med and pSMAD are broadly shared between the two strains. Previous work in cell culture experiments [16] and *Xenopus* embryogenesis (Warmflash, 2012) expressed exogenous Venus-SMAD4, the mammalian homolog of Medea, to measure BMP signaling dynamics. An exogenous GFP-Med was also recently reported by another group [44–45]. However, our tool is the first endogenous reporter of BMP signaling in the *Drosophila* embryo. In the early fly embryo, there is no input from Activin signaling, as BMP ligands are the only expressed TGF-beta family signaling molecules. In tissues that activate the R-SMAD Smox, which also complexes with Medea for nuclear translocation [46–47], nuclear Med concentration would not distinguish between Smox and Mad activity. Attempts to tag Mad, however, do not result in homozygous viable flies. Moreover, in our hands, C-terminal and N-terminal tags of Mad do not show dynamic localization.

### Integration at BMP responsive promoters

Our data reveal that time-integrated BMP signaling is a significantly better predictor of transcriptional activation than instantaneous signaling levels. This finding highlights integration as a key decoding strategy in a developmental context where gene expression changes must be made within minutes. The ability of BMP target genes to respond based on cumulative signaling rather than a simple threshold allows for spatial patterning from small signaling changes via a rapidly evolving gradient, an insight that emerges directly from our high-resolution, quantitative methods in live embryos. This mechanism is complementary to the adaptive response observed in cultured myoblast cells [14,48] and proposed for the control of cell division in fly wing imaginal discs [10]. In adaptive systems, the transcriptional response depends on the relative rate of change of signaling. In these systems, the observed responses to signaling dynamics, as well as the adaptation time [48], are on timescale of hours. Similarly, in an *in vitro* model of hPSC differentiation, where BMP signal integration was also observed, this integration takes place over the course of days and BMP-induced SOX2 expression was proposed to act as an integrating factor over such long time frames [14]. Differently from these previous studies, our work shows that cells in the early *Drosophila* embryo integrate BMP signals on the scale of tens of minutes, and thus likely do not have sufficient time between input and output to show adaptive behavior or rely on the accumulation of a downstream transcriptional target to integrate information.

Our ATAC-seq data, as well as previously published ATAC-seq from carefully staged cycle 13 embryos [49], suggest that the chromatin openness at BMP target genes is not actively changing during the time frame of promoter firing. This suggests that deposition of epigenetic marks that increase accessibility at these promoters does not occur. It is possible that the dwell time of Mad/Medea, Zen or some other cofactor on chromatin is very long and could build up over time. However, most transcription factor-chromatin interactions are short-lived and thus rely more heavily on concentration for activity at the promoter [50]. Thus, we propose a different theoretical rationale for why the time integral of signaling might be predictive of gene expression dynamics.

We have focused on the timing of the initial activation (“first firing”) of BMP target gene promoters, as this appears to be the key regulatory step. Once transcription begins, nuclei typically maintain a transcriptionally active, bursting state for the hour-long period of interphase prior to gastrulation, as shown in our data and previous studies [33,44]. With this in mind and considering that gene expression can be triggered by very low levels of integrated BMP activity, we propose that the onset of gene expression can be described as a first passage time problem [51]. In physics, first passage time problems arise naturally when computing the first time that a system reaches a given configuration [52]. Mathematically, that requires computing over time the probability that a system has not visited that configuration yet. The simplest formulation of this problem is for a Poisson process where the probability rate of reaching the configuration over time is described by *r*(*t*). It can be shown that the first passage time probability density function *P*(*t*) is given by:

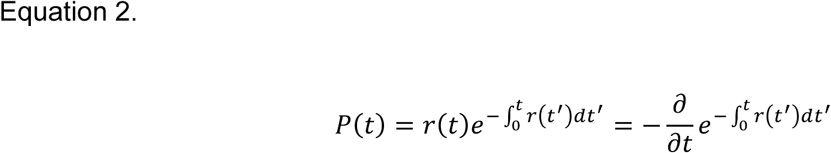

Thus, the time-integral of the rate of gene activation naturally appears in the probability of gene activation over time. Assuming that rate depends on the concentration of Mad/Medea, i.e. *r*(*t*)∼*c*(*t*), where *c*(*t*) indicates the Mad/Meda concentration, we would expect the time of gene activation to be determined by the time integral of the signaling dynamics. This provides a different intuitive explanation for the dependency of gene activation on integrated signaling activity: the longer a cell is exposed to a high signaling rate, the more likely it is to reach the transcriptionally active state. Integration emerges naturally from the statistics of a stochastic activation process: higher or more sustained BMP signaling increases the likelihood of a promoter firing event, and this dependence on both concentration and duration naturally gives rise to an integrated relationship. In this sense, transcriptional activation behaves as if the system is integrating the signal over time, even in the absence of a dedicated molecular integrator.

We propose that this probabilistic explanation of promoter activity is useful for patterning when the decision time is very rapid. With long integration times, specificity of the expression domain would be lost. The cells on the dorsal midline must make these initial cell fate decisions in the matter of tens of minutes before gastrulation begins and the amnioserosa is needed for morphogenesis. In addition, the BMP signaling gradient is zygotically encoded rather than maternally deposited, such that cells do not have a long time or large dynamic range of information in which to produce outputs from BMP signaling inputs. This idea might differ for gene expression patterns on the ventral side of the embryo or along the anterior-posterior axis as these patterns have much more time to evolve. Indeed, it is compatible with the observation that gene expression domains on the dorsal side of the embryo are much more graded than the sharp stripes of gene expression observed on the ventral side and along the anterior-posterior axis of the embryo.

### Zen catalyzes the BMP response at target genes

Our data show that *zen* is essential for the rapid onset of transcription in response to BMP activity. In *zen^7^* mutant embryos, expression of BMP target genes was delayed, resulting in narrower patterns of gene expression. This indicates that Zen lowers the threshold required for transcriptional activation, enabling nuclei to respond to low, integrated BMP signaling inputs. In the absence of Zen, this rapid integration-based decoding mechanism breaks down.

We developed an endogenous LlamaTag-Zen allele, allowing us to track Zen protein dynamics *in* vivo. We found that Zen protein is present across the dorsal side of the embryo early in nuclear cycle 14. This contradicts previous feed-forward models in which BMP signaling reinforces a dorsally restricted Zen gradient to activate target genes. Instead, our data support a model in which early, uniform expression of Zen driven by Zelda [28,53], primes the embryo for BMP responsiveness, enabling transcriptional responses during a narrow developmental time window. Zen may act by promoting transcriptional machinery recruitment or stabilization at target gene promoters. While *zen* has been shown to be required for expression of some BMP target genes, our work suggests that *zen* may be required at most BMP targets for correct spatial-temporal expression. We do not believe Zen acts to affect chromatin accessibility or to directly promote SMAD binding at target gene promoters. Rather, Zen may assist in the recruitment of transcriptional machinery to the same target genes. Future studies should explore whether this competence role depends on early Zen binding at BMP target promoters and how its refinement to the dorsal midline influences subsequent morphogenetic events. More broadly, our findings illustrate how combinatorial regulation by a competence factor (Zen) and a dynamic morphogen (BMP) allows for fast, robust patterning, a mechanism likely relevant to other developmental contexts requiring rapid gene expression in response to transient signals.

### Combinatorial signaling reinforces spatial expression domains

Our data also reveal an important limitation inherent to this integration-based model. In *sog* mutant embryos, where the BMP gradient is expanded and flattened, many lateral cells receive integrated BMP levels comparable to those observed close to the midline in wild-type embryos. Yet, these nuclei fail to activate transcription of BMP target genes at predicted levels. This suggests that integrated signaling alone is not sufficient to determine transcriptional outcomes across the entire dorsal domain. Instead, additional spatially patterned regulatory mechanisms likely restrict the effective range of BMP responsiveness. This highlights the importance of considering not only how much and for how long a signal is present, but also where in the embryo the signal is being interpreted. One possible explanation for the failure of signal integration to predict gene activation in lateral regions is that additional co-factors are required to modulate competence to respond to BMP signaling. Transcriptional repressors, such as *brinker* [28,54–55], might be active at the promoters of these BMP target genes in *sog* mutant embryos.

Altogether, our findings suggest that spatial patterning of gene expression in response to a dynamic morphogen gradient can be quickly and robustly patterned by reading out integrated thresholds. However, integrated signaling thresholds are not the full picture. Competence factors like Zen modulate the sensitivity of target promoters to signaling input, and combinatorial signaling ensures that only cells within a defined spatial domain can respond to low levels of BMP. This layered regulatory logic which combines quantitative integration and spatial gating, may represent a general strategy for rapid but robust pattern formation in early development.

**Supplemental Figure 1: BMP signaling dynamics from fixed tissues**

**A)** Reference curve for developmental staging based on the extent of cellularization. The percentage of cellularization front progression was quantified over time from three wild-type embryos imaged under brightfield with 1-minute time resolution. Data were fit with a smoothing spline. This cellularization front curve was used to stage additional fixed embryos during nuclear cycle 14 with ∼10-minute temporal resolution.

**B–D)** Heatmaps showing spatially binned nuclear pSMAD levels over time along the dorsal-ventral axis in wild-type (B), *sog* mutant (C), and *sog* heterozygous (D) embryos. Each row represents the average pSMAD signal in 5 μm-wide spatial bins over time.

**Supplemental Figure 2: GFP-Med is a robust reporter of BMP signaling activity**

**A)** Comparison of nuclear GFP-Medea and pSMAD staining in fixed, staged embryos. Nuclear signal intensities were averaged in space (5 μm bins) and time, and color-coded by developmental stage (light to dark blue, t = 0–60 minutes of nuclear cycle 14). GFP-Medea levels closely track pSMAD levels over time and space.

**B)** Overlap of ChIP-seq peaks for GFP-Med and pSMAD in wild-type embryos. Peaks were called from experiments using anti-GFP (in GFP-Med embryos), with control anti-GFP ChIP in wild-type used to filter non-specific peaks. The normalized anti-GFP (from GFP-Med embryos) or anti-pSMAD (from wild type embryos) ChIP signal is plotted over these peak locations.

**C–D)** Examples of ChIP-seq signal for GFP-Med and pSMAD at representative BMP target gene loci, showing strong concordance in peak location and intensity.

**E)** Immunofluorescence image of a late cycle 14 embryo from a GFP-Mad heterozygous female, co-stained for pSMAD.

**F)** Quantification of nuclear GFP-Mad and pSMAD intensities across the dorsal-ventral axis for the embryo shown in (E), demonstrating that the GFP-Mad does not recapitulate the pSMAD gradient as effectively as GFP-Med.

**G)** Time-resolved dynamics of nuclear GFP-Medea accumulation in *sog* heterozygous embryos (n=2). Data are shown as average nuclear intensity in 5 µm spatial bins from the dorsal midline, with 1-minute temporal resolution.

**H)** Rate of change in GFP-Medea levels across the dorsal-ventral axis in *sog* heterozygous embryos.

**Supplemental Figure 3. Additional characterization of transcriptional dynamics in WT and *sog* mutant embryos.**

**A–C)** Single-cell traces of nascent transcriptional activity in live embryos expressing MS2 reporters for *ush* (A), *doc2* (B), and *c15* (C), showing that once a gene is activated, nuclei generally remain transcriptionally active until gastrulation (black lines) and less than 10% of nuclei that fire the promoter turn off (red lines).

**D–F)** Temporal expression profiles for *c15* (D, n=36), *doc2* (F, n=24), and *tup* (H, n=37) in *sog* mutant embryos from fixed time-course smFISH data. Dotted lines indicate regions with >10% of nuclei on in wild type embryos.

**G–I)** Same as (D–F), but in *sog* heterozygous embryos (n=18, 24, and 24 respectively). Dotted lines indicate regions with >10% of nuclei on in wild type embryos.

## Experimental Procedures

### *Drosophila* Stocks and Husbandry

A list of all fly strains used in this study can be found in the Key Resources Table. *Drosophila melanogaster* flies were grown and maintained at room temperature on standard molasses food (Archon, Cat #B20101). Fly strains and crosses for imaging were maintained at 25°C for at least 24 hours before embryo collection. For FISH experiments, flies were placed in an egg lay chamber with apple juice agar plates and yeast paste for 2 hours. The plates were removed, and embryos were aged for an additional 2 hours at 25°C before fixation. For live imaging experiments, female flies were allowed to lay eggs for 2-4 hours at 25°C before collection.

### Generation of Transgenic *Drosophila*

All transgenic lines in this study were generated by CRISPR/Cas9 genome editing with homologous recombination. Guide RNA (gRNA) sequences were inserted into pCFD3-dU6:3 (Addgene #49410) by BbsI digestion and T4-mediated ligation. To create the donor plasmid for C-terminal tagging of *Mad* or *Med*, either eGFP or TagRFP was cloned into pBluescript downstream of the start codon of the target gene with 1kb of flanking genomic DNA on either side. 3xP3-dsRed flanked by loxP sites was added between the 5’ homology arm and the start codon for screening transformants. Notably, *Med* is expressed as two different isoforms, one maternally deposited and the other zygotically transcribed according to modENCODE Development RNAseq data (FlyBase). In this study, we targeted the maternally expressed isoform (*Mad-RA*), which disrupts the first exon of the zygotic isoform. The *Med* N-terminal was the same in all isoforms and maternally expressed. Donor plasmids for MS2 reporters of *c15* and *doc2* live transcription were generated by cloning 24 MS2 stem loops within the 5’UTR of the endogenous gene locus, flanked by 1kb of homology arms in the MCS of pBluescript. The 3xP3-dsRed flanked with loxP was again cloned between the 5’ homology arm and the MS2 stem loops. gRNA and donor plasmids for homology repair were injected into embryos of Cas9 expressing flies (BDSC #51323) by Model Injection Systems or BestGene. The dsRed construct was used for screening and then excised by crossing to CreW expressing balancer flies for at least one generation. A similar method was also used to target the N-terminus of *zen* with a GFP LlamaTag.

### smFISH Probe Preparation

Probes were designed using Oligostan [56]. 20-48 smFISH probes per gene were either ordered as oligopools from IDT or mixed in the lab in EB to yield a combined final concentration of 0.833 µM for each probe. To anneal FLAP-Y or FLAP-X probes to the probe set, 2uL of the probe set was mixed with 1uL of the appropriate fluorescently labeled FLAP oligo in 10uL of 1X NEB 3 buffer without BSA (10 mM NaCl, 5 mM Tris-HCl, 1 mM MgCl2). FLAP probes were annealed in the thermocycler using the following program: 85°C for 3 minutes, 65°C for 3 minutes, 25°C for 5 minutes. Probes were stored away from light at 4°C for no more than one week before use.

### Immunofluorescence and smFISH

Embryos were collected from apple juice agar plates and dechorionated in 50% Clorox (4% bleach) for 1 minute followed by thorough rinsing with water and drying on a paper towel. Embryos were then transferred to a scintillation vial containing 3 mL of freshly prepared 4% formaldehyde in PBS (or 10% formalin) and 3 mL of heptane to form a biphasic mixture. Embryos were fixed by vigorous shaking on a tabletop shaker for 20 minutes. After fixation, the lower aqueous phase was carefully removed, and 5 mL of methanol was added. The vials were shaken by hand for 30–45 seconds to facilitate vitelline membrane removal. Embryos were transferred to 1.5 mL microcentrifuge tubes using a glass pipette and washed three times with 500 µL methanol. To retain GFP fluorescence when fixing GFP-Med embryos, these methanol washes were omitted and instead, the methanol was quickly replaced with ethanol following vitelline membrane removal.

Following fixation, embryos were rehydrated through a methanol/PBT (PBS + 0.1% Triton X-100) gradient with sequential 5-minute washes in 3:1, 1:1, and 1:3 methanol:PBT, followed by two 5-minute washes in 100% PBT. Embryos were incubated with pSMAD-1/5 primary antibody diluted 1:200 in PBT overnight at 4 °C. After primary antibody incubation, embryos were washed three times for 5 minutes each in PBT. Secondary antibody incubation was performed in PBT for 2 hours at room temperature, protected from light, using a 1:500 dilution. This was followed by three 5-minute washes in PBT, also protected from light. Embryos were then incubated with DAPI (1:50,000 in PBT) for 10 minutes, followed by a final 5-minute wash in PBT. Stained embryos were mounted in Vectashield mounting medium for imaging.

For smFISH immunostaining experiments, prior to the addition of DAPI, embryos were subjected to a second round of 4% formaldehyde fixation for 20 minutes at room temperature on a rotator. Embryos were washed twice for 2 minutes in 15% formamide in 1X SSC. For hybridization, two mixes were prepared. Mix 1: 12.5 µL 20X SSC, 2.5 µL 20 mg/mL BSA, 4.25 µL 50 µg/µL tRNA, 37.5 µL 100% formamide, 5 µL FLAP probes, and 65.75 µL water. Mix 2: 2.5 µL RNase inhibitor, 66.25 µL 40% dextran sulfate, and 53.75 µL water. If both a FLAPX and FLAPY probe were used, 5uL of each probe set was added to Mix 2 and the water was reduced by 5uL. Mix 1 and Mix 2 were combined and used to hybridize embryos for 72 hours at 37°C on a nutator in the dark. Following hybridization, embryos were washed twice for 5 minutes at 37°C in fresh 15% formamide in 1X SSC, and twice more for 5 minutes in PBT at room temperature. For nuclear staining, embryos were incubated with DAPI (1:50,000 in PBT) for 20 minutes, followed by three 5-minute washes in PBT. Embryos were then mounted in Vectashield on microscope slides for imaging.

### *Drosophila* Genetics and Genotyping

To capture live movies of BMP signaling activity and MS2 reporter activity, HisiRFP, MCP-mCherry, and GFP-Med must all be maternally deposited. Due to these genetic constraints, only *ush*-MS2 reporter flies could be imaged concurrently with GFP-Med by crossing *ush*-MS2;GFP-Med flies with HisiRFP;MCP-mCherry and collecting females to back-cross to *ush*-MS2;GFP-Med males. MCP-mCherry constructs on the second chromosome that would allow imaging of third-chromosome MS2 reporters alongside GFP-Med were too easily photobleached by even low-level laser power to successfully complete these experiments. Instead, HisRFP;MCP-GFP females were crossed to males carrying third chromosome MS2 reporters (*c15* and doc2). We observed Zen protein concentration and localization in embryos maternally loaded with GFP (vasa>GFP) and a nuclear Histone marker (HisRFP) and expressing Zen-LlamaTag from the endogenous locus.

To capture live GFP-Med signal in mutant embryos, we generated GFP-Med,*zen^7^* recombinant chromosomes. For both *zen* and *sog* mutants, we imaged several embryos laid by heterozygous females. The genotype of each embryo was determined based on the final shape of the BMP gradient and if embryos were able to successfully complete dorsal closure.

To genotype fixed embryos stained with smFISH, we used a second smFISH probe. For *zen* mutants, we used a FLAP-X probe against *gal4*, which was expressed from the balancer chromosome. The number of transcription spots could discriminate between one or two copies of the balancer chromosome. For *sog* mutants, we used a FLAP-X probe against *sog* itself. This method could not distinguish between male and heterozygous embryos, so we carefully sorted embryos based on the width of the pSMAD staining, though this method could only be used for embryos in later time points when the shape of the gradient is more distinct from wild type and *sog* mutants.

### Confocal Imaging

Fixed tissue confocal imaging was performed on a Leica SP8 and live imaging on a Stellaris confocal microscope using a 20×/0.75 numerical aperture oil-immersion objective at 2.5x zoom. Images were acquired at 1,024 x 1,024 resolution and z-stacks were acquired with 1.5 μm resolution. Brightfield imaging of the cellularization front was obtained using transmitted light from the argon laser for fixed tissue data by focusing on the midplane of the embryo at 2x zoom. For live imaging, z-stacks were acquired with 1 minute time resolution. Some MS2-MCP movies were obtained with 20 second time resolution.

### Single Embryo ATAC-seq

ATAC-seq libraries were prepared from single *Drosophila* embryos staged by observation under a dissection microscope. Individual embryos were arrayed in halocarbon on apple juice agar plates and monitored for the onset of cellularization. When an embryo reached early nuclear cycle 14, a timer was started. 15 minutes later, the embryo was dechorionated in 4% bleach for 1 minute, rinsed in water, and transferred to the inverted cap of a 1.5 mL LoBind tube containing 10 µL of ATAC lysis buffer (10 mM Tris pH 7.4, 10 mM NaCl, 3 mM MgCl₂, 0.1% Igepal CA-630). The embryo was lysed using a microcapillary pestle. The lysate was centrifuged at 500 × g for 10 minutes at 4 °C, and the nuclear pellet was flash frozen on dry ice.

Nuclear pellets were thawed on ice and resuspended in 10 µL of tagmentation reaction mix (5 µL 2× Tn5 Buffer, 2.5 µL Tn5 enzyme, and 2.5 µL water; Illumina). Samples were incubated at 37 °C for 30 minutes with shaking at 800 rpm. DNA was purified using the MinElute Reaction Cleanup Kit (Qiagen) and eluted in 22.5 µL of Buffer EB.

Libraries were amplified using NEBNext Q5 Master Mix and dual-indexed primers. The optimal number of PCR cycles was determined by a qPCR side reaction using SYBR Green, and the main PCR was carried out accordingly. Amplified libraries were cleaned using Ampure XP beads, eluted in 15 µL EB, and quantified using the Qubit High Sensitivity DNA Assay. Library size distribution was assessed on an Agilent Tapestation using D5000 reagents.

### Chromatin Immunoprecipitation

Chromatin immunoprecipitation followed by Tn5-mediated library preparation (Tn5-ChIP-seq) was performed using batches of ∼100 staged *Drosophila* embryos per replicate. Embryos were collected on yeasted apple juice plates and dechorionated in fresh 4% bleach for 1 minute, followed by extensive rinsing in distilled water and PBS with 0.5% Triton X-100. Embryos were fixed in a biphasic mixture of 6 mL heptane and 2 mL PBS/0.5% Triton X-100 containing 180 µL of 20% paraformaldehyde. Fixation proceeded for 15 minutes at room temperature with vortexing (30 seconds) and shaking at 120 rpm. The reaction was quenched by addition of 1 mL PBS/0.5% Triton X-100 with 125 mM glycine. Embryos were washed three times in ice-cold PBS/0.5% Triton X-100 and genotyped using a dissection microscope.

To select homozygous mutant embryos, the *zen^7^*,GFP-Med recombinant chromosome was balanced over a TM3 balancer containing a halo rescue construct. This was crossed to a second chromosome deficiency line that removes the *halo* locus. Therefore, screening for embryos with the halo phenotype allowed for selection of homozygous mutants. We confirmed in later sequencing results that about 90% of the reads we obtained did come from mutant chromosomes, so the halo rescue construct is not 100% penetrant. These embryos and wild type controls were flash frozen in liquid nitrogen and stored at −80°C for at least 12 hours.

Fixed frozen embryos were resuspended in RIPA buffer (1× RIPA supplemented with 1 mM DTT and protease inhibitors). Lysates were sonicated (Branson Digital Sonifier, ¼” microtip, 20% output, 15 seconds, four rounds) to release chromatin. Immunoprecipitation was performed overnight at 4 °C with primary antibody and pre-blocked Protein G Dynabeads (blocked with 3% BSA and 0.2 mg/mL yeast RNA in PBS/0.1% Triton X-100).

Following immunoprecipitation, chromatin-bound beads were washed twice in tagmentation wash buffer (10 mM Tris pH 7.5, 5 mM MgCl₂) and incubated for 40 minutes at 37 °C in 21 µL of tagmentation mix (10 µL water, 10 µL 2× Nextera Tagmentation Buffer, 1 µL Tn5 transposase). After tagmentation, samples were washed sequentially in standard ChIP wash buffers (low-salt, high-salt, and TE), and eluted in 100 µL elution buffer (50 mM Tris-HCl pH 8.0, 10 mM EDTA, 1% SDS) with 5 µL of 20 mg/mL Proteinase K. Crosslink reversal was carried out for 4 hours at 65 °C with shaking.

Eluted DNA was purified using the Qiagen MinElute Reaction Cleanup Kit and PCR-amplified using NEBNext Q5 Master Mix with unique dual-indexed primers. The optimal number of amplification cycles was determined using a qPCR side reaction with SYBR Green. Final libraries were cleaned with 1.8× Ampure XP beads and eluted in 15 µL of Buffer EB. Libraries were quantified using Qubit and assessed for fragment size on an Agilent Tapestation with D5000 reagents.

### Fixed Tissue Staging

Fixed *Drosophila* embryos were staged based on the progression of the cellularization front, as observed in brightfield microscopy. Embryos were imaged in lateral orientation using transmitted light, and their developmental stage was assessed by comparing the position of the cellularization front to a reference time-lapse movie of embryos imaged live. The live cellularization front reference embryo expressed HisiRFP, so all times could be calculated relative to anaphase of nuclear cycle 13. To convert cellularization progression into elapsed time, the percentage of cellularization at each time point in the live movie was quantified and used to generate a smooth curve describing cellularization front progression over time. Each fixed embryo’s degree of cellularization was manually measured in Fiji, placed on the fitted curve, and used to infer the corresponding time since the start of cellularization. Embryos were then assigned to temporal bins as follows: 0–15 minutes, 15–30 minutes, and subsequent 10-minute intervals up to 60 minutes.

### Nuclear Segmentation and Tracking

Fixed and live *Drosophila* embryo images were processed to segment nuclei and RNA signals using a combination of CellProfiler and custom MATLAB scripts. Nuclear segmentation was performed in CellProfiler using either DAPI staining (for fixed samples) or fluorescent histone markers (for live imaging). For each embryo, nuclear objects were identified in 2D across z-planes, and custom MATLAB scripts were used to reconstruct 3D nuclear positions by stacking across z and, for time-lapse data, tracking nuclei over time.

MS2 transcriptional foci and smiFISH-labeled RNA dots were also segmented in CellProfiler using standard spot-detection pipelines. CellProfiler was also used to pair nuclei with their daughter MS2 dot(s) by Parent-Child detection. First, nuclei were expanded until touching, creating nuclear neighborhoods, and dots within those neighborhoods were then paired with the corresponding nuclear object. Output measurements from CellProfiler including nuclear positions, dot counts, and fluorescence intensities were imported into MATLAB for downstream analysis.

### Quantification and Normalization of GFP-Med

GFP-Med levels were quantified in segmented nuclei from live *Drosophila* embryos. Nuclear segmentation was performed as described above, and mean nuclear GFP fluorescence intensity was extracted for each nucleus.

To correct for experiment-to-experiment variability in fluorescence intensity due to technical factors such as laser power and detector sensitivity, a two-step normalization procedure was applied. First, background GFP-Med levels were estimated for each embryo by calculating the average nuclear fluorescence in lateral regions at least 100 µm from the embryonic midline, where signaling is minimal. This background value was subtracted from all nuclear intensities within the same embryo to correct for nonspecific signal and baseline differences.

Second, to account for embryo-to-embryo variability, fluorescence values were scaled using a reference embryo of the appropriate genotype imaged under standard conditions. This was done by minimizing the sum of squared differences between binned GFP-Med intensity profiles from the experimental and reference embryos over a defined region up to 45 microns from the midline. All nuclear intensities in the experimental dataset were then multiplied by this scale factor to enable quantitative comparisons across embryos and imaging sessions.

Integrated signaling was defined as the cumulative integral of GFP-Med over time for each spatial bin, calculated by fitting a curve to temporal signal and integrating the fitted function from 20 minutes after anaphase of cycle 13 up to each time point. We chose the 20 minute timepoint due to the observation that some nuclear GFP-Med signal remains for about 20 minutes after nuclear envelope breakdown and reformation of nuclear cycle 13. We are unsure if this nuclear signal represents active signaling, but it is relatively uniform across the dorsal side of the embryo and would therefore affect all positions equally.

### Midline Determination

To quantify spatial position relative to the dorsal midline, different approaches were used for live imaging and fixed embryos. In live MS2 datasets, the dorsal midline was calculated at a late time point (55 minutes into cellularization). The midline was defined as the peak of nuclear GFP-Medea signal along the dorsal-ventral axis at this time point. Once determined, the midline position was held constant, and nuclei in earlier frames were tracked backward in time to maintain their spatial relationship relative to the midline. In fixed embryos, where each image represents a single time point, the midline was identified independently for each embryo. A smoothing spline was fit to the pSMAD fluorescence intensity profile across the dorsal-ventral axis. To avoid distortions from signal variability at the flanks, the curve was fit using only the central region of the profile, excluding the extreme lateral portions of the embryo. The position of maximum pSMAD intensity within this central region was taken as the dorsal midline.

### Mapping Target Gene Expression to Spatial-Temporal Domains

To analyze the relationship between RNA expression and GFP-Med, nuclei were grouped into spatial bins based on their distance from the dorsal midline. For both MS2 live imaging and smFISH fixed data, the embryo divided into 3 µm-wide bins extending up to 60 µm from the midline. Within each bin, RNA expression was quantified as either the presence or absence of MS2 transcriptional spots (live imaging) or smFISH puncta (fixed embryos) in each nucleus. In fixed samples, the fraction of nuclei with detectable RNA signal (i.e., the probability of expression) was computed within each spatial bin. To infer RNA expression dynamics at higher temporal resolution, a logistic curve was fit to the expression probabilities across all fixed embryos and bins. This model provided a smooth estimate of the probability of gene expression as a function of time since the start of cellularization, allowing inference of expression at minute-by-minute resolution. Each nucleus in live MS2 movies or fixed smFISH datasets was matched to its corresponding spatial bin and assigned a developmental time point based on cellularization staging (described above) or time since anaphase of nuclear cycle 13. For each bin and time point, RNA expression levels were then compared to GFP-Med fluorescence in corresponding spatial-temporal bins in wild-type or mutant embryos to assess the correlation between signaling strength and transcriptional output. These relationships were modeled in Matlab using the Hill Equation, and fit parameters extracted.

### Quantification of Zen-LlamaTag

Zen-LlamaTag fluorescence was quantified in fixed embryos to assess protein localization and dynamics. Nuclear segmentation was performed as described above, and cytoplasmic regions were defined by expanding the nuclear masks by a fixed pixel radius and subtracting the original nuclear region, thereby creating a perinuclear cytoplasmic shell for each nucleus. Both nuclear and cytoplasmic fluorescence intensities were extracted from the raw Zen-LlamaTag channel. To correct for photobleaching and technical variability across the imaging field or between embryos, the nuclear-to-cytoplasmic (NC) ratio of Zen-LlamaTag signal was calculated for each nucleus. This ratiometric approach enabled internal normalization within each embryo and eliminated the need for external background measurements. The NC ratio was used for downstream comparisons of Zen-LlamaTag dynamics across time points.

### ATAC-seq and ChIP-seq Data Analysis

Dual-indexed ChIP-seq or ATAC-seq libraries were sequenced as 150 bp paired-end reads on an Illumina NovaSeq X platform (Admera Health). Raw FASTQ files were demultiplexed and adapter sequences were trimmed using TrimGalore. Cleaned reads were aligned to the *Drosophila melanogaster* dm6 reference genome using Bowtie2. Aligned reads were sorted and duplicates were flagged using Picard MarkDuplicates. For downstream analysis, only uniquely mapped reads with a mapping quality score of at least 10 were retained. For ATAC-seq data, embryos were genotyped using a custom SNP caller in R. For ChIP-seq and ATAC-seq data, replicates were merged and imported into R and converted to GenomicRanges objects for manipulation. Genome-wide coverage was calculated in 10 bp bins, and normalized to library size to generate counts-per-million (CPM) coverage tracks. These tracks were visualized using the GViz package in R.

For ATAC-seq data, peaks of accessible chromatin were identified using MACS. Peaks were called for wild-type and mutant samples using merged replicates for each genotype. Comparative analyses between wild-type and mutant embryos were performed by overlapping peak sets and assessing differential peak presence.

For ChIP-seq datasets, GFP-Med peaks were called in wild type embryos using MACS with a control input derived from a GFP ChIP performed in a no-GFP background. This control accounted for non-specific enrichment and background signal. Resulting peak sets were filtered based on q-value and p-value. Enrichment at peaks in *zen^7^* mutant embryos was visualized using the GViz package in R. All data is available at GEO accession number GSE298384.

## Acknowledgements

We thank members of the Di Talia Lab, Dr. Brigid Hogan, and Dr. Bernard Mathey-Prevot for scientific discussion and feedback on the manuscript. This work was supported by the following grants from the National Institutes of Health: 5R01-GM136763 to S.D., 1F32GM145070-01 to S.E.B., and 5K99 HD115781 to S.E.B. This work was also supported by ab Innovation in Stem Cell Science Award from the Shipley Foundation, Inc. to S.D.

