## Supplementary figures and images for "Rapid transcriptional response to a dynamic morphogen by time integration"

### Supplemental Figure 1

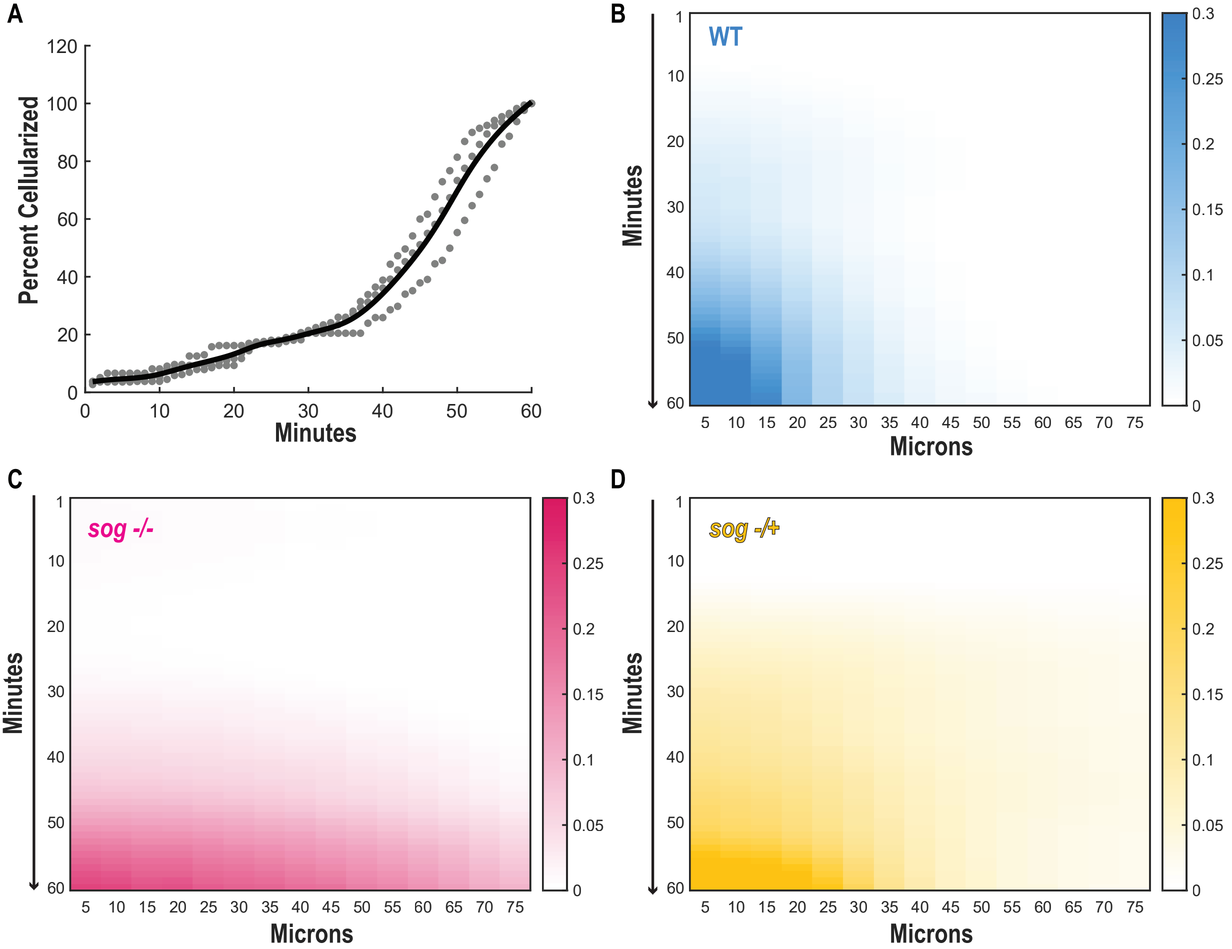

### Supplemental Figure 2

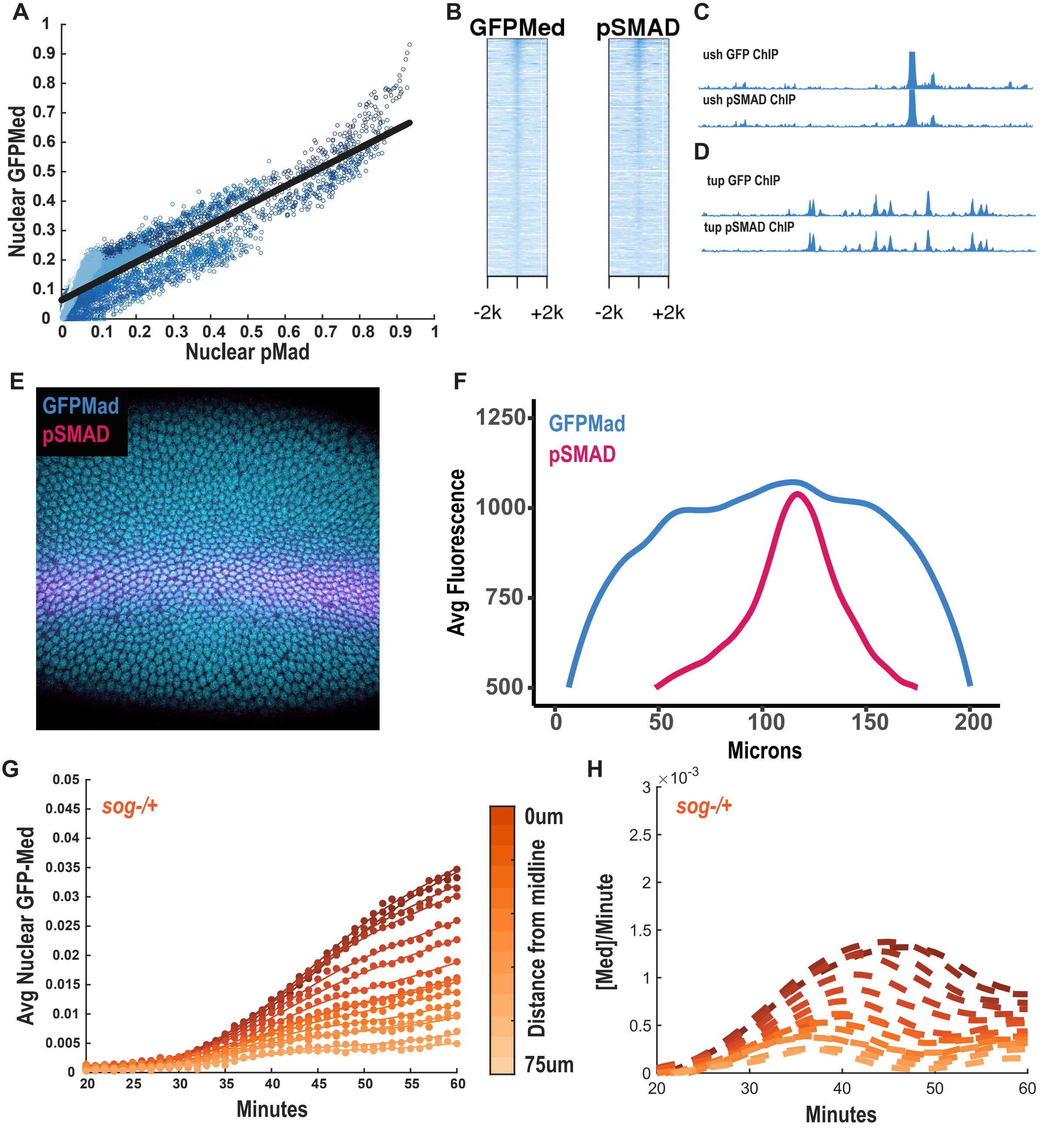

### Supplemental Figure 3

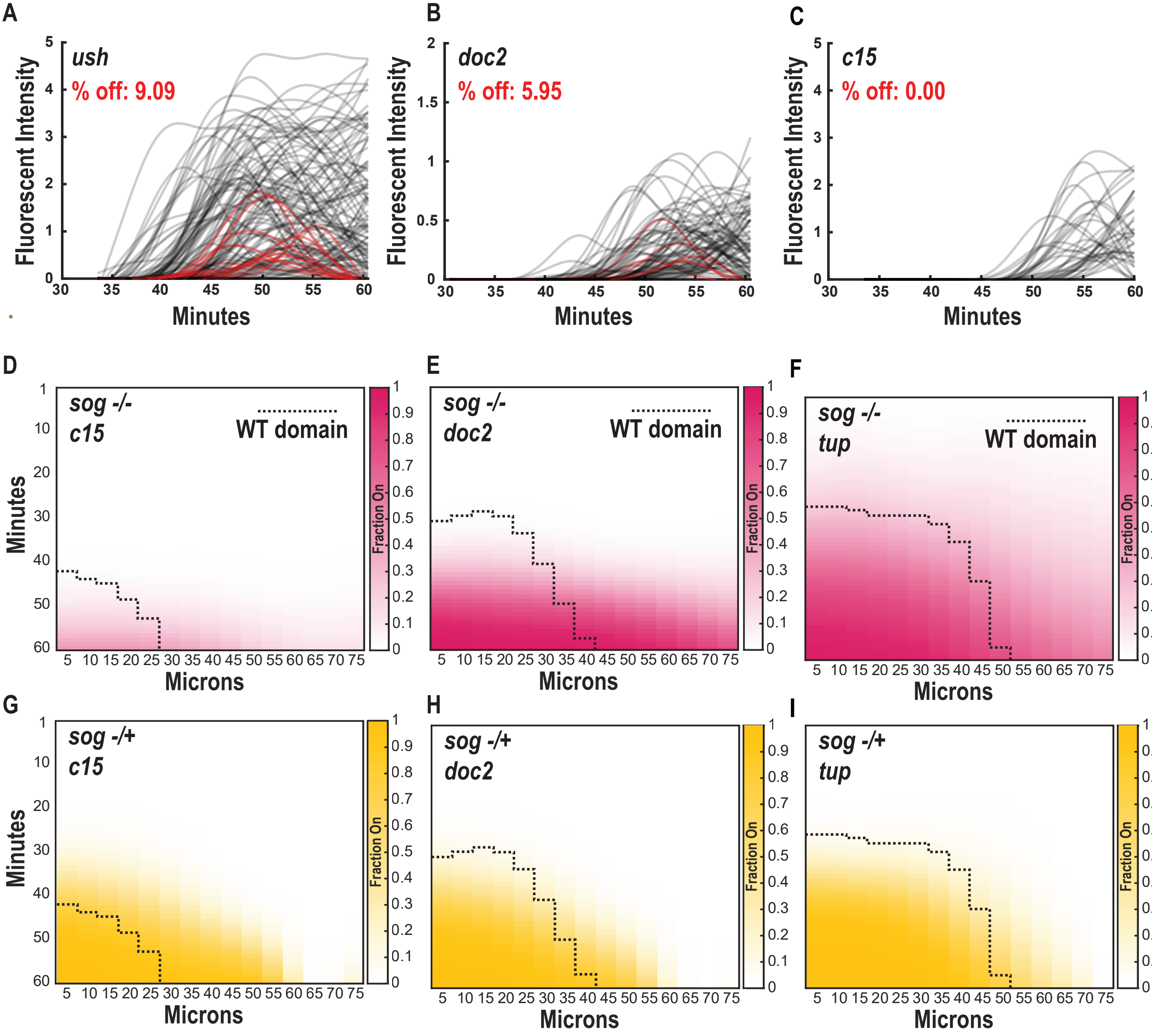
